# *Eurotium cristatum*, a new fungi probiotic from Fuzhuan brick tea, alleviated obesity in mice by modulating gut microbiota

**DOI:** 10.1101/518068

**Authors:** Dingding Kang, Meng Su, Yanwen Duan, Yong Huang

## Abstract

Obesity is one of the major public health problems worldwide, mainly resulting from unhealthy lifestyles and diet. Gut microbiota dysbiosis may also lead to obese humans and animals. Modulating gut bacteria through fecal transplantation, the use of probiotics or certain dietary supplements, could normalize gut microbiota and subsequently alleviate obesity. Daily consumption of Fuzhuan brick tea (FBT) or its extracts have been observed to alleviate obesity in humans and experimental animals. In this study, high-fat diet-induced dysbiosis of gut microbiota in C57BL/6J mice was partially reversed by consumption of *Eumtium cristatum*, the dominant fungi during the manufacturing and storage of FBT. *E. cristatum* was able to modulate both gut fungi and bacteria composition, based on the analysis of microbiota composition of mice fecal samples. *E. cristatum* increased acetate and butyrate-producing bacteria in mice gut, and produced five times more butyrate than both obese and normal mice. Our results suggested that *E. cristatum* may be used as a fungi probiotic to beneficially modulate gut microbiota and to alleviate obesity in humans.

## Introduction

Obesity is a disease associated with numerous health problems, mainly caused by a shift in lifestyles towards less physical expenditure and more hyper-caloric foods^1^. It is closely linked with chronic and low-grade inflammation, which may cause insulin resistance, type 2 diabetes, fatty liver disease or cardiovascular disease^2–4^. Based on two recent population analysis from 1975-2016 or 1980-2015, about 10% of the world population, including over 100 million young people, were obese^5^. Obesity epidemic becomes a major threat to public health in modern human history^6^. Obesity treatment may include bariatric surgery or changing of lifestyles to increase exercise, which reduce calorie intake^7^. Anti-obesity drugs, such as phentermine or orlistat, could suppress appetite or inhibit the absorption and utilization of fat^8^. Alternatively, regulating intestinal microbes has recently been used for the treatment of obesity, because microbiota dysbiosis plays prominent roles in the development of obesity^9–12^. A variety of probiotics, either individually or more often used as cocktails, were shown to alleviate obesity in a high-fat diet (HFD) fed rodents. For example, both *Bifidobacterium* and *Lactobacillus* were shown to attenuate HFD-induced obesity in animals and humans, including weight loss, reduced visceral fat and improved glucose tolerance^13–15^. Most of these bacteria produce short-chain fatty acids (SCFAs), such as acetate and butyrate, which reduce gut luminal pH and maintain an acidic environment^16,17^. Previous studies also highlighted the importance of SCFAs to improve chronic inflammatory diseases and to promote colonocyte health^18^. SCFAs were reported to suppress production of pro-inflammatory cytokines IL-1β, enhance IL-10 expression and activate regulatory T cells(T_reg_), to maintain intestinal homeostasis^19^.

Humans have prepared and consumed a variety of fermentation products, such as pickles, sauerkraut, sourdough bread, artisanal cheeses, yogurts and kefir, craft beers and kombucha (fermented tea), in which various bacteria and fungi species are present, dated back to the seventh millennium before Christ^20,21^. Fermentation is very useful to preserve food and increase their flavor and nutrition^22^. Fuzhuan brick tea (FBT, also named Fu brick tea) is a fermented tea in the presence of a dominant fungi species named *Eurotium cristatum*, which is yellow-colored and is thus also called “golden flora” 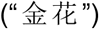 in Chinese^23^. Human consumption of FBT dated back to the 16^th^ century^24^, while recent pilot dietary interventions on humans in China have also shown that FBT consumption could improve metabolic disease conditions, which were summarized in Supplementary Table 1. FBT consumption or polysaccharides extracted from FBT were shown to reduce obesity, hyperlipidemia and arterial stiffening, as well as to improve metabolic syndrome in animals^25–28^. The water or ethanol extracts of FBT reduced obesity through normalizing gut microbiota in mice^25,29^.

The dominant fungi *E. cristatum* in FBT play an important role in the flavor, color and health benefits of FBT^30^. We hypothesized that live *E. cristatum* in FBT may survive in the gastrointestinal tract during tea consumption. It might have served as a fungi probiotic to modulate gut microbiota, which led to the observed anti-obesity effects of FBT. In this study, we evaluated the effects of dietary supplementation of live *E. cristatum* to prevent diet-induced obesity in mice. Our study supports the use of *E. cristatum* as a fungi probiotic to modulate gut microbiota against obesity, which has been used by humans for about 600 years.

## Results

### *E. cristatum* may survive in mouse intestine

In order to clarify if *E. cristatum* could survive during tea brewing and consumption, 10^4^ spores of *E. cristatum* isolated from FBT, were heated in water containing 20% (v/v) glycerol at 50, 70, 80, 85, 90 and 100 °C for 2, 5 or 10 min, respectively (Supplemental Fig.1). *E. cristatum* survived even after 10 min in 90 °C and 2 min in 100 °C. *E. cristatum* was next mixed with drinking water and fed to Kunming mice. *E. cristatum* could be isolated from mouse feces, even without feeding of *E. cristatum* for two weeks (Supplemental Fig. 2). There were no significant differences in body weight among mice fed with or without *E. cristatum*, as well as mice fed with the water extracts or filtered water extracts of Fuzhuan brick tea. These results suggested that *E. cristatum* from Fuzhuan brick tea is non-toxic and could survive in mice intestine.

### *E. cristatum* reduced HFD-induced obesity in C57BL/6J mice

The HFD-induced C57BL/6J obese mouse model was used to explore if live *E. cristatum* may have any anti-obesity effect, in comparison to mice that received normal chow diet (NCD). The high-fat diet mice that received FBT or filtered Fuzhuan brick tea (FFBT), in which live *E. cristatum* were removed by filtering the FBT extracts using a 0.22 μm filter, were also used as controls (Fig. 1a-e). The mice in HFD group showed significant increases in body and liver weights, and epididymal fat accumulation, compared with mice in the NCD group (Fig. 1a-e). In contrast, the high-fat diet mice that received *E. cristatum* (10^3^ CFU/day) in drinking water showed significantly decreased body weight gain, liver weight and epididymal fat accumulation. The FFBT-treated mice also showed significant decrease in liver and epididymal fat, while FBT-treated mice only had statistically-insignificant lower body weight, in comparison to HFD-fed mice.

**Fig. 1.**
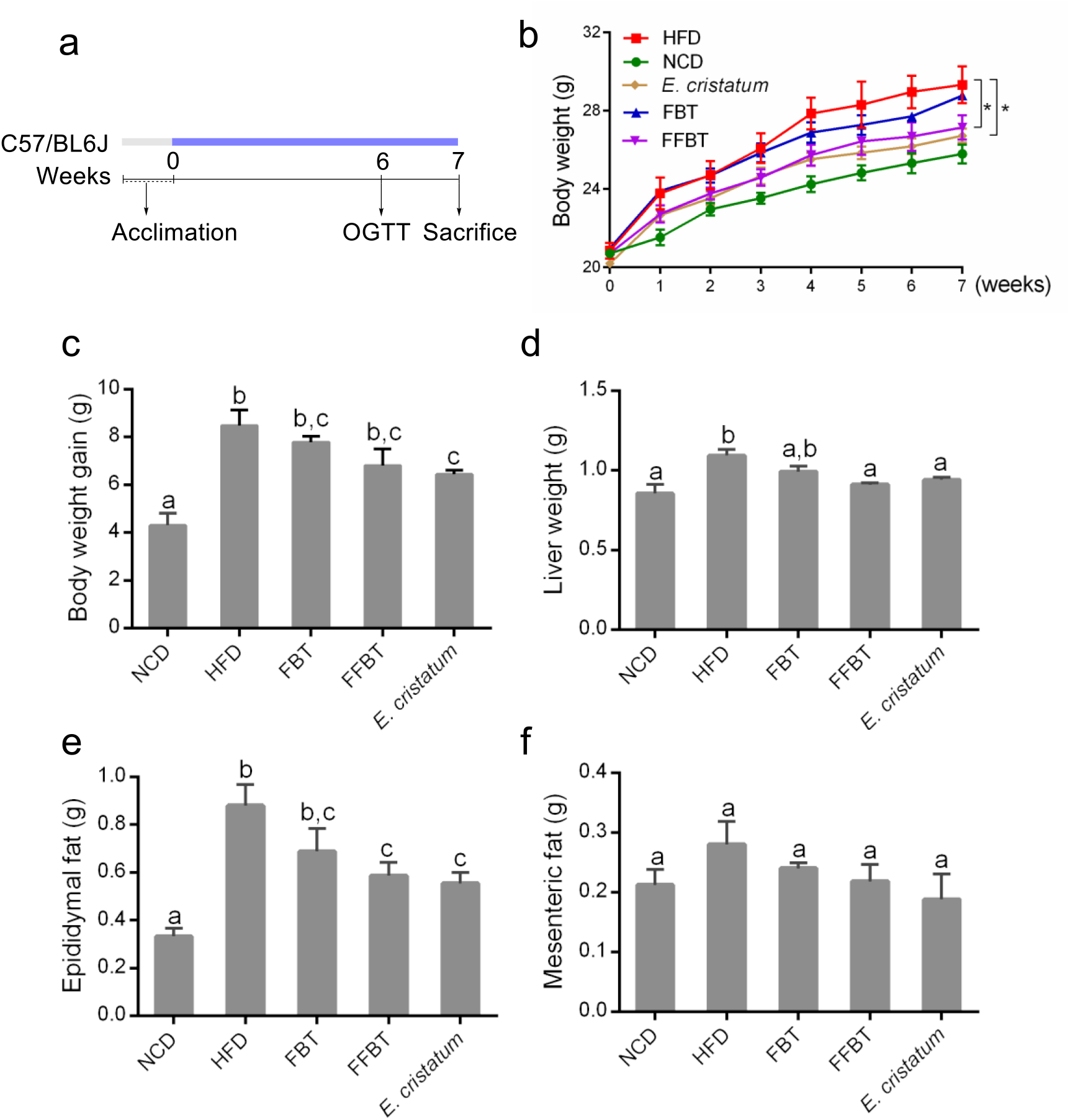
Dietary *E. cristatum* reduced body weight and fat accumulation in HFD-induced mice. **a**, The animal protocol. **b**, Effects of *E. cristatum*, FBT and FFBT on body weight in 7 weeks. **c**, Body weight gain. **d**, Liver weight. **e**, Epididymal and mesenteric fat. **f**, Data are expressed as mean ± s.e.m. Body weight differences in **b** were analyzed using unpaired two-tailed Student’s *t*-test (\**P*<0.05). Graph bars in figure **c, d, e** and **f** marked with different letters on top represent statistically significant results (*P*<0.05) based on Newman–Keuls *post hoc* one–way ANOVA analysis, whereas bars labeled with the same letter correspond to results that show no statistically significant differences. In the case, whereas two letters are present on top of the bar, each letter should be compared separately with the letters of other bars to determine whether the results show statistically significant differences.

### *E. cristatum* increased glucose tolerance in HFD-fed mice

The effects of *E. cristatum*, FBT or FFBT consumption on glucose homeostasis of HFD-fed mice were next determined (Fig. 2a-c). Compared to HFD-fed mice that received only drinking water, the HFD-fed mice with daily consumption of *E. cristatum*, FBT or FFBT had reduced fasting blood glucose levels (FBG) and increased glucose tolerance. Since endotoxemia controls the production of pro-inflammatory cytokines in target tissues and may lead to chronic inflammation in HFD-fed mice^31^, the serum lipopolysaccharide (LPS) levels in these mice were further examined (Fig. 2d). Both FBT and FFBT consumption reduced the LPS level in HFD-fed mice, while *E. cristatum* had no significant effect.

**Fig. 2.**
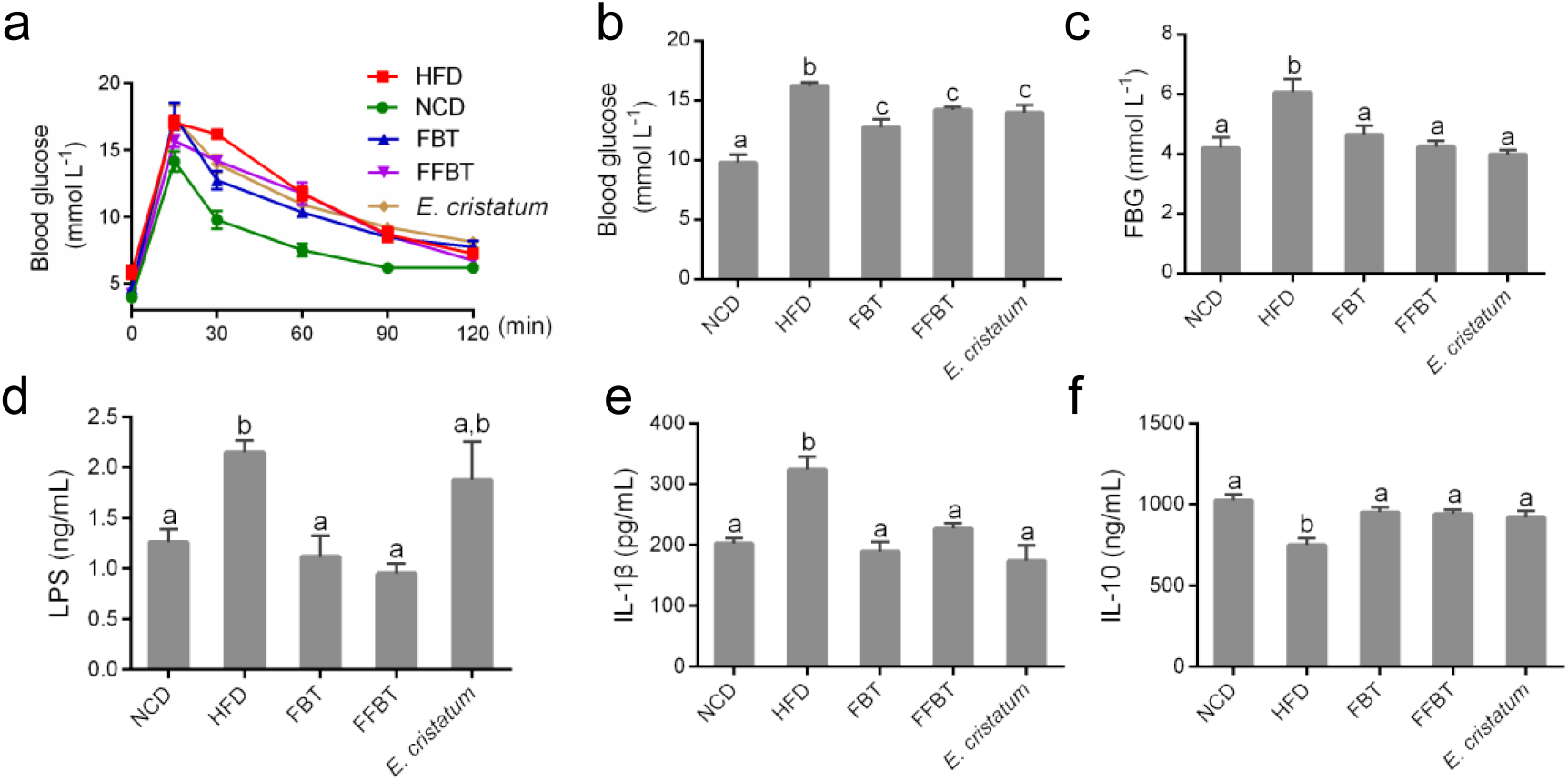
*E. cristatum*, FBT or FFBT reduced HFD-induced metabolic endotoxemia and systemic chronic low-grade inflammation, and improved glucose homeostasis. **a**, Effects of *E. cristatum*, FBT and FFBT on blood glucose using OGTT test. **b**, Blood glucose of 30 min of OGTT. **c**, Fasting blood glucose (FBG). **d**, LPS level. **e**, IL-1β. **f**, IL-10 in hepatic tissues. Data are expressed as mean ± s.e.m. Graph bars in figure **b, c, d, e** and **f** marked with different letters on top represent statistically significant results (*P*<0.05) based on Newman-Keuls *post hoc* one-way ANOVA analysis.

### *E. cristatum* reduced inflammation in HFD-fed mice

HFD-induced obese mice have been observed to produce higher levels of pro-inflammatory cytokines in hepatic tissues, including interleukin-1-β (IL-1β) and interleukin-6 (IL-6), while the level of the anti-inflammatory cytokine interleukin (IL-10) was reduced in obese animals^32^. The IL-1β and IL-10 in hepatic tissues of the sacrificed mice were thus measured by enzyme-linked immune sorbent assay (ELISA) (Fig. 2e-f). The HFD-fed mice produced about 300 pg/mL IL-1β, while the consumption of *E. cristatum*, FBT or FFBT reduced the IL-1β level to about 200 pg/mL, similar to the NCD-fed mice. Similarly, the levels of anti-inflammatory cytokine IL-10 were also restored to those in NCD-mice through consumption of *E. cristatum*, FBT or FFBT.

### *E. cristatum* altered the fungi diversity in mice gut

Since consumption of *E. cristatum* alone had significant anti-obesity effects of the HFD-fed mice (Fig. 1-2), we hypothesized that *E. cristatum* may be able to modulate mice gut microbiota through its interaction with commensal fungi and bacteria^33,34^. Therefore the composition, abundance and function of mice gut microbiota were next analyzed by high-throughput sequencing of the internal transcribed spacer 2 (ITS2) and 16S rRNA of caecal stool samples of the above C57BL/6J mice. After removing unqualified sequences, a total of 741,205 raw reads and an average of 37,060 ± 23,543 reads per mouse sample were obtained for ITS2 rDNA regions. Rarefaction of chao1 and observed OTUs indicated that the sequencing depth covered most of the fungi diversity, including rare new phylotypes (Supplemental Fig. 4a-b).

To evaluate fungi diversity in mice gut, the fungi β diversity was measured. Constrained principal-coordinate analysis (CPCoA) based on the similarity index revealed distinct clustering of fungi composition for each mice group (Fig. 3a). *E. cristatum-* or FBT-treated HFD-fed mice were closer to NCD-fed mice. Further hierarchical clustering analysis did not reveal distinct separation of each mice group. Fungi taxonomic profiling in the genus level of intestinal fungi revealed that *Euroteomycetes* were more abundant in *E. cristatum-*treated mice than those of other groups, which suggested that *E. cristatum* had survived in mice intestine (Fig. 3c-d). Statistical analysis of metagenomic profiles (STAMP) of gut fungi also revealed several OTUs belonging to *Scolecobasidium, Eurotium, Penicillium* and *Cosmospora*, which were significantly altered in different mice groups (Fig. 3e-f).

**Fig. 3.**
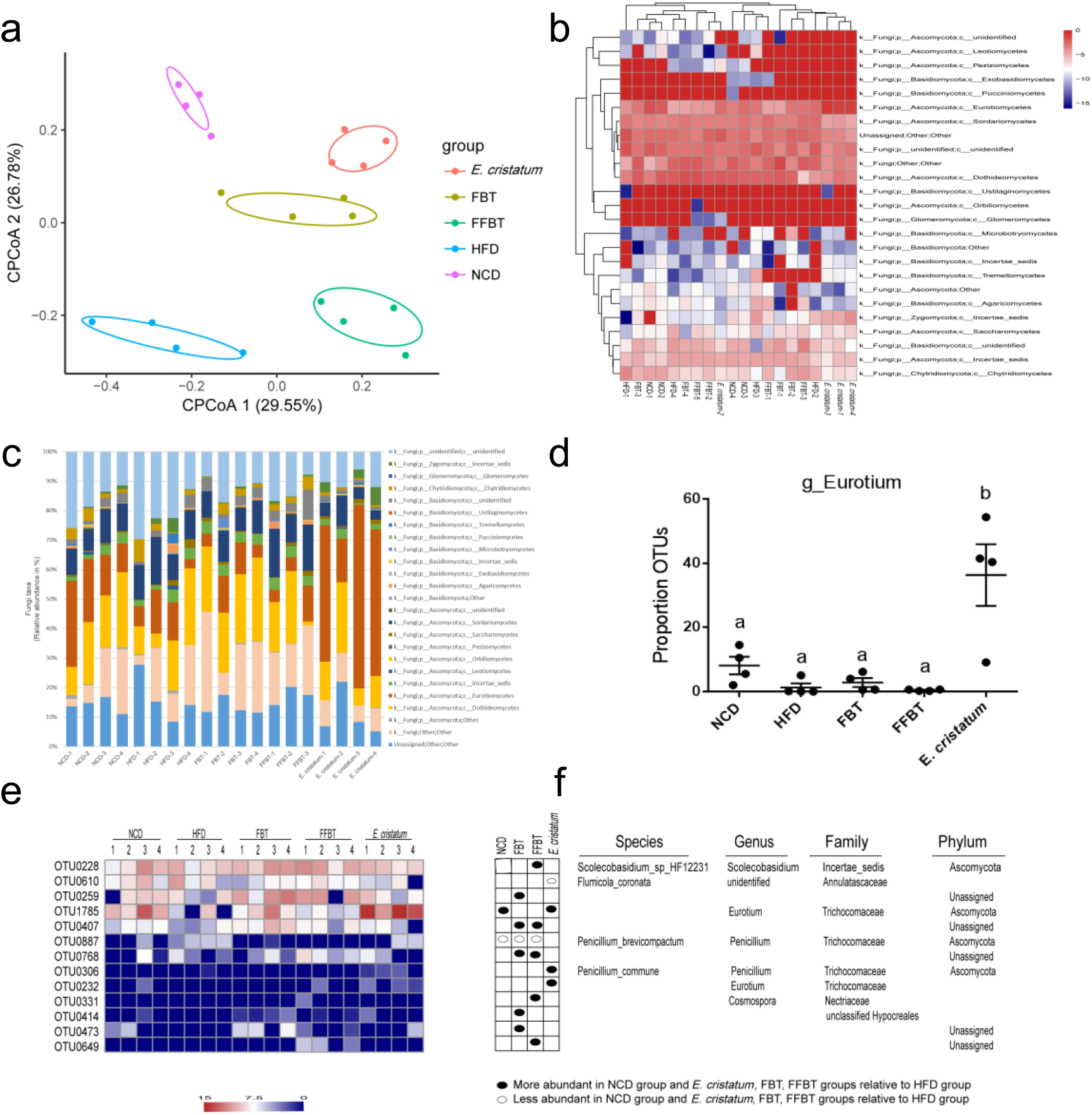
Dietary *E. cristatum*, FBT and FFBT beneficially altered the gut fungi composition. **a**, Constrained principal-coordinate analysis based on the cao similarity index with the PERMANOVA significance test of the ITS2 region. **b**, Hierarchical clustering showing the relative abundance of representative OTUs. **c**, Fungi taxonomic profiling in the family level of intestinal fungi from different mice groups. **d**, The relative abundances of *E. cristatum*. **e**, The abundance of OTUs significantly altered by *E. cristatum*, FBT and FFBT in HFD-fed mice based on STAMP. **f**, Represented bacterial taxa information (species, genus, family, and phylum) of OTUs from e.

### Dietary *E. cristatum* beneficially altered the gut bacteria

The V4 region of 16S rRNA of mice gut bacteria were also obtained, which included a total of 765,760 raw reads and an average of 30,630±17,252 reads per mouse sample. The rarefaction curves analysis of chao1 and observed OTUs revealed that most of the commensal bacteria diversity were covered (Supplemental Fig. 4). The β diversity of the gut bacteria was also evaluated using CPCoA-based Bray–Curtis similarity index, which revealed distinct clustering of bacteria composition for each mice group (Fig. 4a). Compared with other groups, *E. cristatum-*treated HFD-fed mice were closer to NCD-fed mice in both primary and secondary ordination axes. The inter-sample correlation calculated by Pearson coefficient analysis revealed that the gut bacteria of NCD and *E. cristatum* groups were more closely related (Fig. 4b).

**Fig. 4.**
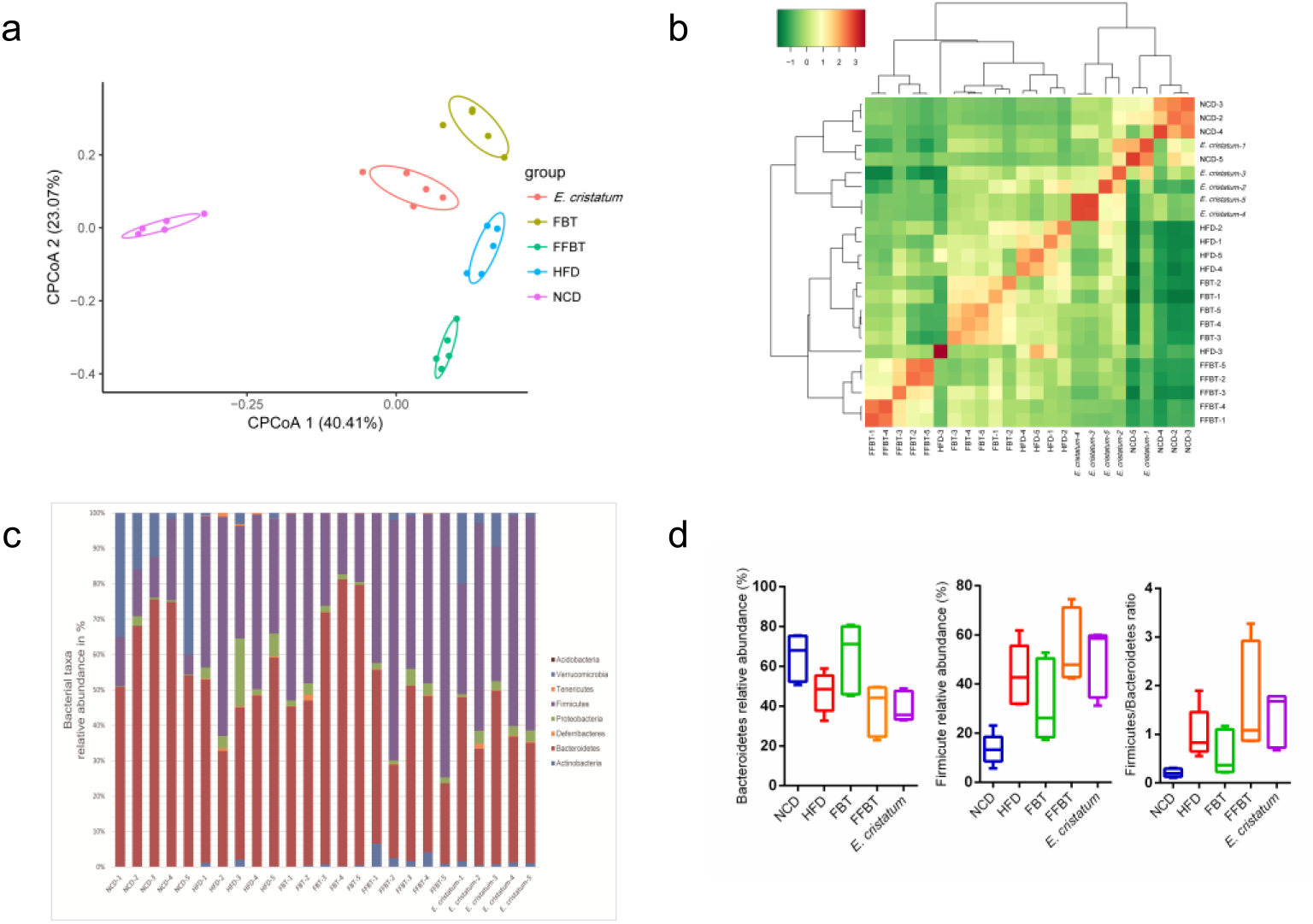
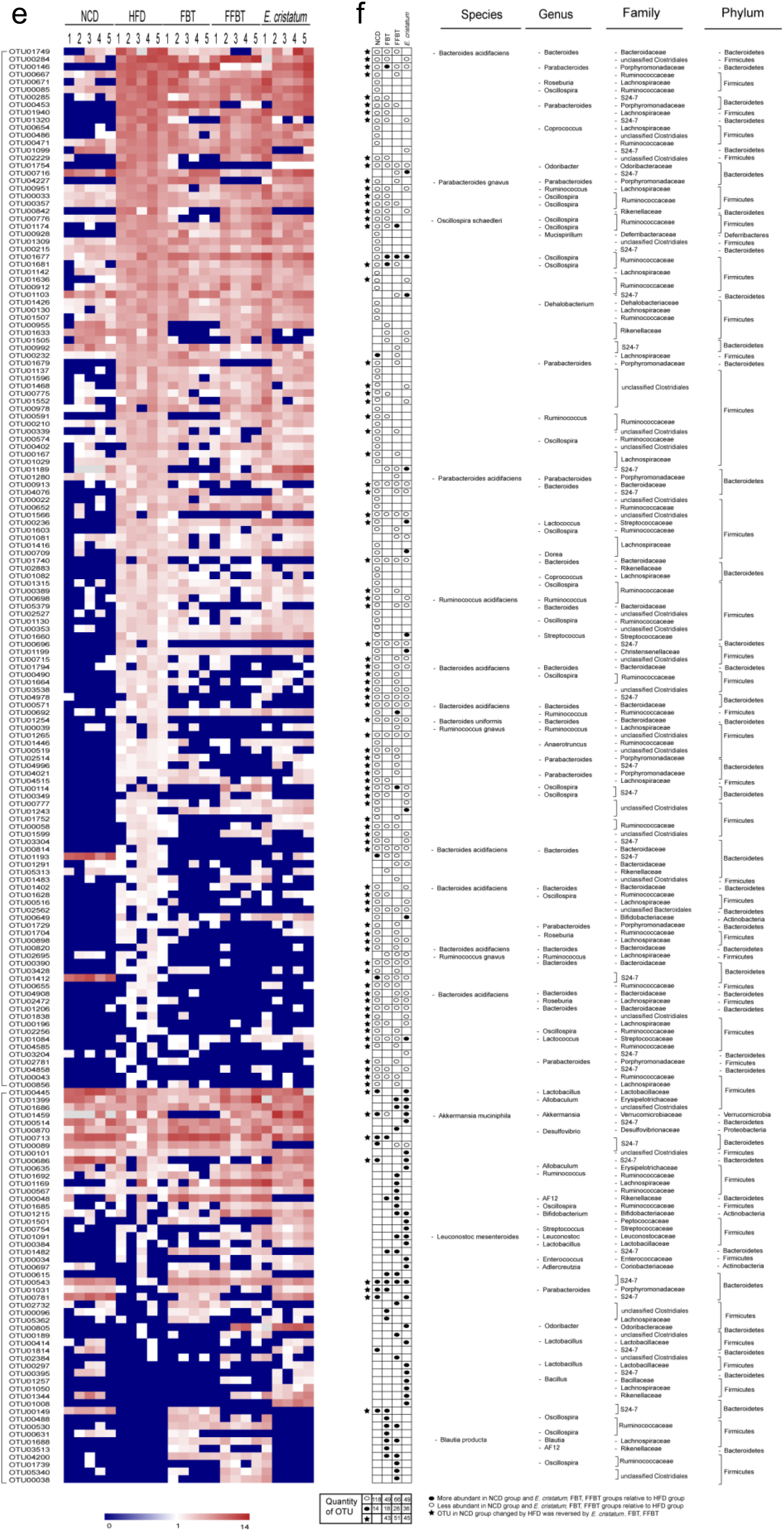
Dietary *E. cristatum*, FBT and FFBT beneficially altered the gut bacterial composition. **a**, Constrained Principal-coordinate analysis based on the Bray–Curtis similarity index with the PERMANOVA significance test about 16S rRNA. **b**, Inter-sample correlation calculated by the Pearson coefficient. **c**, Bacterial taxonomic profiling in the phylum level of intestinal bacteria from different mice groups. **d**, The relative abundances of Bacteroidetes, Firmicutes, and F/B. **e**, The abundance of OTUs significantly altered by *E. cristatum*, FBT and FFBT in HFD-fed mice based on STAMP. **f**, Represented bacterial taxa information (species, genus, family, and phylum) of OTUs from e.

Bacterial taxonomic profiling in the phylum level of intestinal bacteria from different mice groups were next analyzed (Fig. 4c). The gut microbiota of HFD-fed mice were characterized by an increased Firmicutes-to-Bacteroidetes ratio (F/B), which was consistent the previous reports^35,36^. Notably, the treatment of FBT in HFD-fed mice reversed the F/B ratio, while FFBT and *E. cristatum* intervention exhibited limited effects on the relative abundances of Firmicutes and Bacteroidetes and the ratio of F/B (Fig. 4d). STAMP was used to identify the specific bacterial phylotypes of each mouse. Compared with NCD-fed mice, there were 132 bacterial OTUs were significantly altered in HFD-fed mice, in which 118 OTUs increased and 14 OTUs decreased. For HFD-fed mice treated with FBT, FFBT or *E. cristatum*, there were 67 (18 increased and 49 decreased), 96 (26 increased and 66 decreased), 85 (36 increased and 49 decreased) altered OTUs, respectively, compared with HFD-fed mice. Interestingly, 43, 51 and 45 bacterial OTUs in groups of *E. cristatum*, FBT and FFBT, were altered towards NCD-fed mice, respectively. Analysis of the gut bacteria altered by *E. cristatum* consumption indicated that SCFAs-producing bacteria *Lactobacillus* increased over 14 folds, in comparison to HFD-fed mice (Fig. 4e). *Bifidobacterium*, the other SCFAs-producing bacteria, increased dramatically. In contrast, *Odoribacteracae^37^, Parabacterioide*^12^, *Sutterella*^38^ *and Clostridium*, decreased in HFD-fed mice, which were negatively correlated with diet-induced obesity. These results suggested that *E. cristatum*, as well as FBT and FFBT, may modulate the gut bacteria in HFD-fed mice and resulted in a microbiota composition close to that of NCD-fed mice.

### PICRUSt analysis of the potential function of gut microbiota in different groups

In order to study the potential function of gut microbiota in different groups, linear discriminant analysis effect size (LEfSe) was used to infer the relative abundance of Kyoto Encyclopedia of Genes and Genomes (KEGG) pathways, which were predicted by phylogenetic reconstruction of unobserved states (PICRUSt)^39^. The HFD-fed mice had a reduced capacity for energy metabolism and increased capacity of carbohydrate metabolism, consistent with a recent metagenomics-based study^35^. Notably, both galactose metabolism and starch metabolism were significantly lower in *E. cristatum*-treated mice than in those HFD-fed mice. In addition, the energy metabolic pathways, including fatty acid and nucleotide metabolisms were significantly lower in HFD-fed mice and they could be reversed by *E. cristatum* consumption. These metabolic changes could be also identified in the FBT and FFBT groups (Supplemental Fig. 5a-c).

### *E. cristatum* increased SCFAs production in mice gut

The SCFAs analysis of mice fecal samples revealed dramatic differences among these mice groups (Fig. 5). Comparing to NCD-fed mice, the HFD-fed mice showed decreased levels of acetate and propionate in their feces samples, while they had a comparable level of butyrate production. In contrast, consumption of *E. cristatum* in HFD-fed mice significantly increased the amount of acetate and butyrate. Notably, the amount of butyrate in the *E. cristatum* group increased over five times over both NCD-fed and HFD-fed mice. The amount of butyrate in the mice fecal samples in FBT mice group also increased significantly. The increasing amount of acetate and butyrate in *E. cristatum-treated* mice was consistent with the increased acetate- and butyrate-producing bacteria in mice gut, such as *Bifidobacterium* and *Lactobacillus* (Fig. 3-4).

**Fig. 5.**
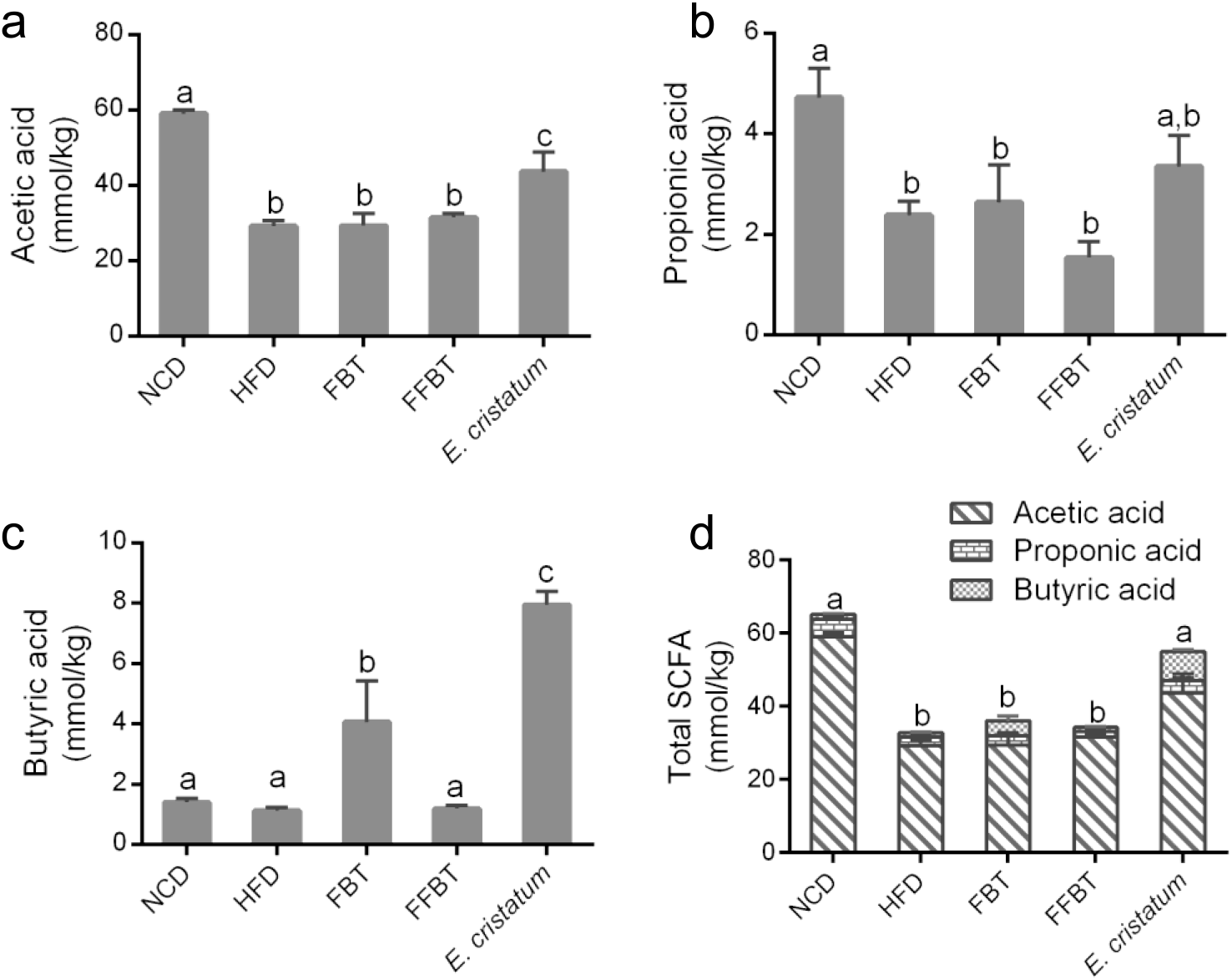
Effects of *E. cristatum*, FBT and FFBT on SCFAs production in mice gut. **a**, Effects of *E. cristatum*, FBT and FFBT on acetic acid, **b**, Propionic acid, **c**, Butyric acid and **d**, Total SCFAs. Data are expressed as mean ± s.e.m. Graph bars in **a, b, c** and **d** marked with different letters on top represent statistically significant results (*P*<0.05) based on Newman–Keuls *post hoc* one–way ANOVA analysis.

### Determination of the optimal dosage of *E. cristatum* to reduce HFD-induced obesity in C57BL/6J mice

Three different dosages of *E. cristatum*, 10^4^ CFU/day, 10^3^ CFU/day and 10^2^ CFU/day, were employed to HFD-induced C57BL/6J mice for 8 weeks. Alternatively, *E. cristatum* (10^3^ CFU/day) were given to HFD-induced mice only for 3 and 5 weeks, respectively. Consumption of different dosages or various periods of *E. cristatum*, all decreased the body weight gain, liver weight and fat accumulation in HFD-fed mice (Fig. 6). The HFD-fed mice receiving *E. cristatum* (10^3^ CFU/day) for 8 weeks gained slightly less weights than other groups. The *E. cristatum* (10^3^ CFU/day, 8 weeks) group also had better blood glucose tolerance than other mice groups, while higher or lower dosage of *E. cristatum* (10^2^ or 10^4^ CFU/day, 8 weeks) were less effective to improve blood glucose homeostasis of HFD-fed mice, indicated by the OGTT test (Fig. 7). Consumption of *E. cristatum* for 3 or 5 weeks also increased glucose tolerance in HFD-fed mice, which suggested that short-term intervention of the microbiota dysbiosis of HFD-fed mice using *E. cristatum* might also be effective (Fig. 7c-d).

**Fig. 6.**
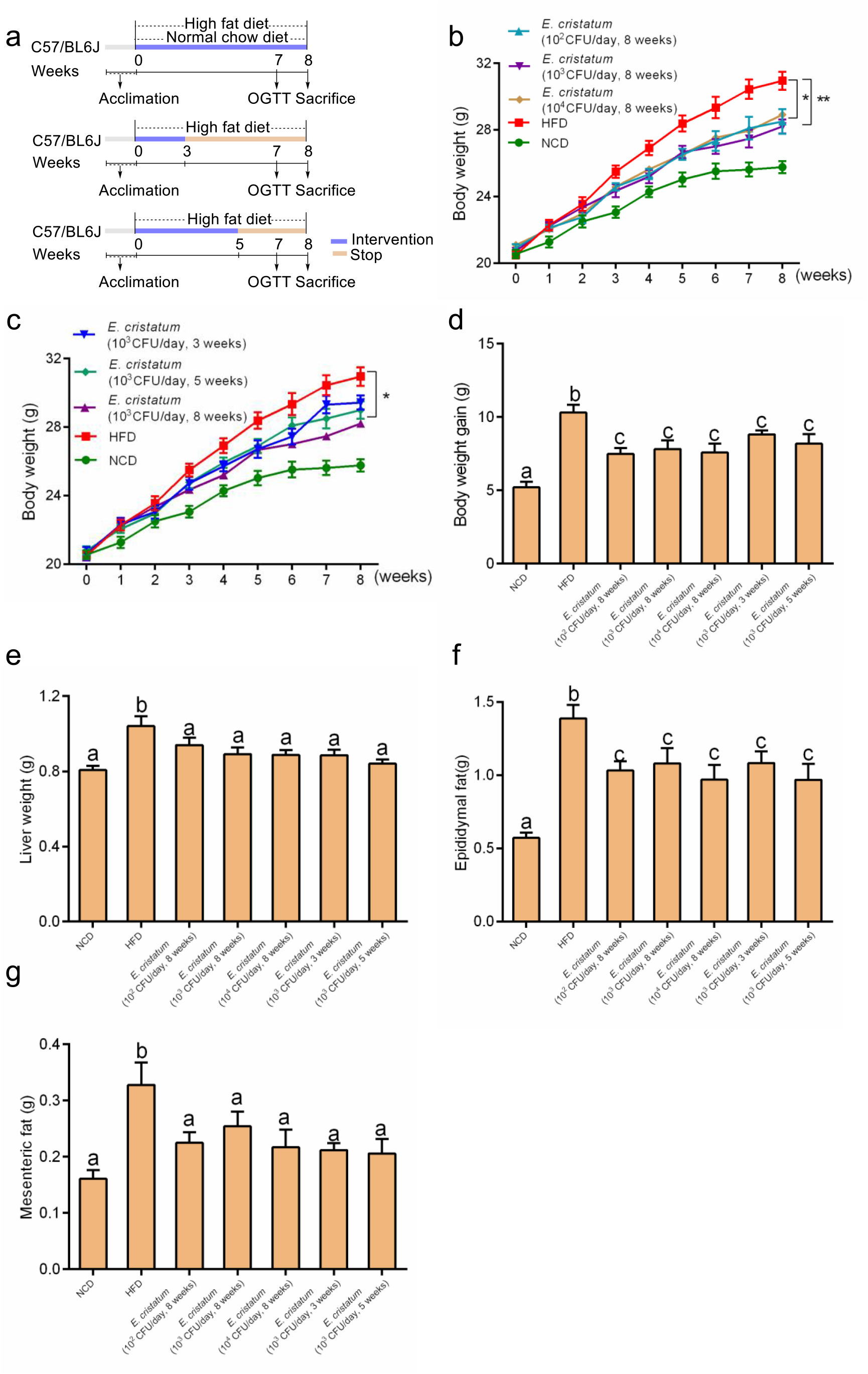
Dietary *E. cristatum* reduced body weight and fat accumulation in HFD-induced mice. **a**, Animal protocol. **b**, Effects of different dosage of *E. cristatum* on body weight of eight weeks. **c**, Effects of consumption various time of *E. cristatum* on body weight of eight weeks. **d**, Body weight gain. **e**, Liver weight. **f**, Epididymal fat. **g**, Mesenteric fat. Data are expressed as mean ± s.e.m. Body weight differences in **b, c** were analyzed using unpaired two-tailed Student’s *t*-test (\**P*<0.05, \*\**P*<0.01). Graph bars in figure **d, e, f** and **g** marked with different letters on top represent statistically significant results (*P*<0.05) based on Newman-Keuls *post hoc* one-way ANOVA analysis.

**Fig. 7.**
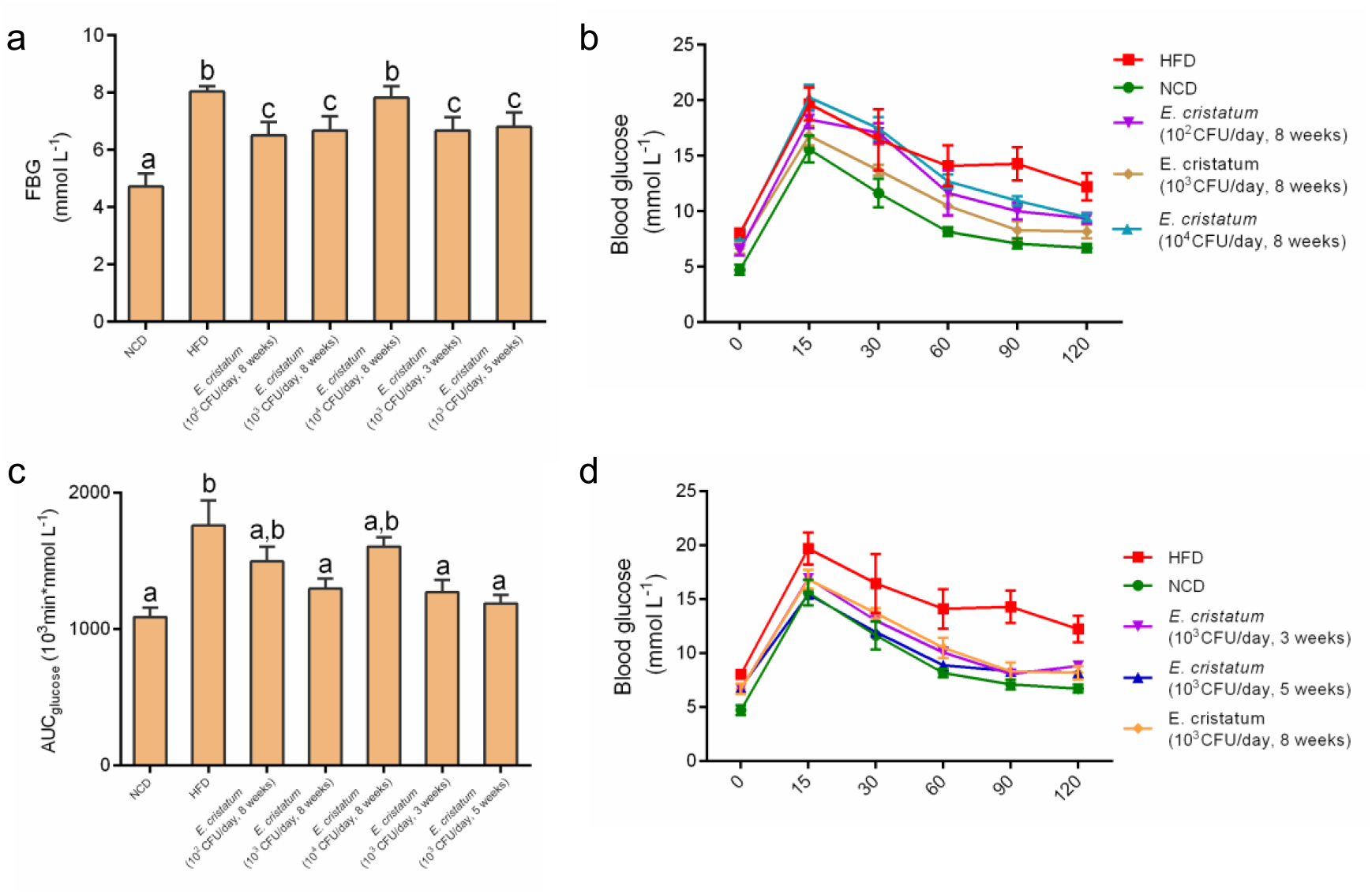
Dietary *E. cristatum* improved glucose homeostasis. **a**, Fasting blood glucose. **b**, Effects of different dosage of *E. cristatum* on blood glucose of OGTT. **c**, Glucose area under the curve (AUCglucose) during OGTT. **d**, Effects of consumption various time of *E. cristatum* on blood glucose of OGTT. Data are expressed as means ± s.e.m. Graph bars in **a** and **c** marked with different letters on top represent statistically significant results (*P*<0.05) based on Newman-Keuls *post hoc* one-way ANOVA analysis.

## Discussion

One of the hallmarks of obesity is the dysbiosis of gut microbiota, and normalization of the gut microbiota through fecal transplantation, consumption of dietary fibers and consumption of certain probiotics may be instrumental to manage the obesity epidemic in the world. In the current study, we hypothesized that *E. cristatum*, derived from Fuzhuan brick tea, may be adopted as a promising fungi probiotic against obesity. In the HFD-fed mice model, *E. cristatum* was shown to alleviate HFD-induced mice obesity, by reducing inflammation and improving their glucose homeostasis. The normalization of gut microbiota and the increased production of butyrate in *E. cristatum*-treated obese mice contributed to the observed anti-obesity effect (Fig. 8). Fuzhuan brick tea extracts and polysaccharides have previously been shown to alleviate HFD-induced mice obesity, probably also through modulation of gut microbiota^25,26,29^. Our study is consistent with these previous studies, in which the FFBT-treated, HFD-fed mice showed reduced liver weight and epididymal fat, and both FBT- and FFBT-treated mice groups showed improved glucose homeostasis and attenuated inflammation, compared to HFD-fed mice.

**Fig. 8.**
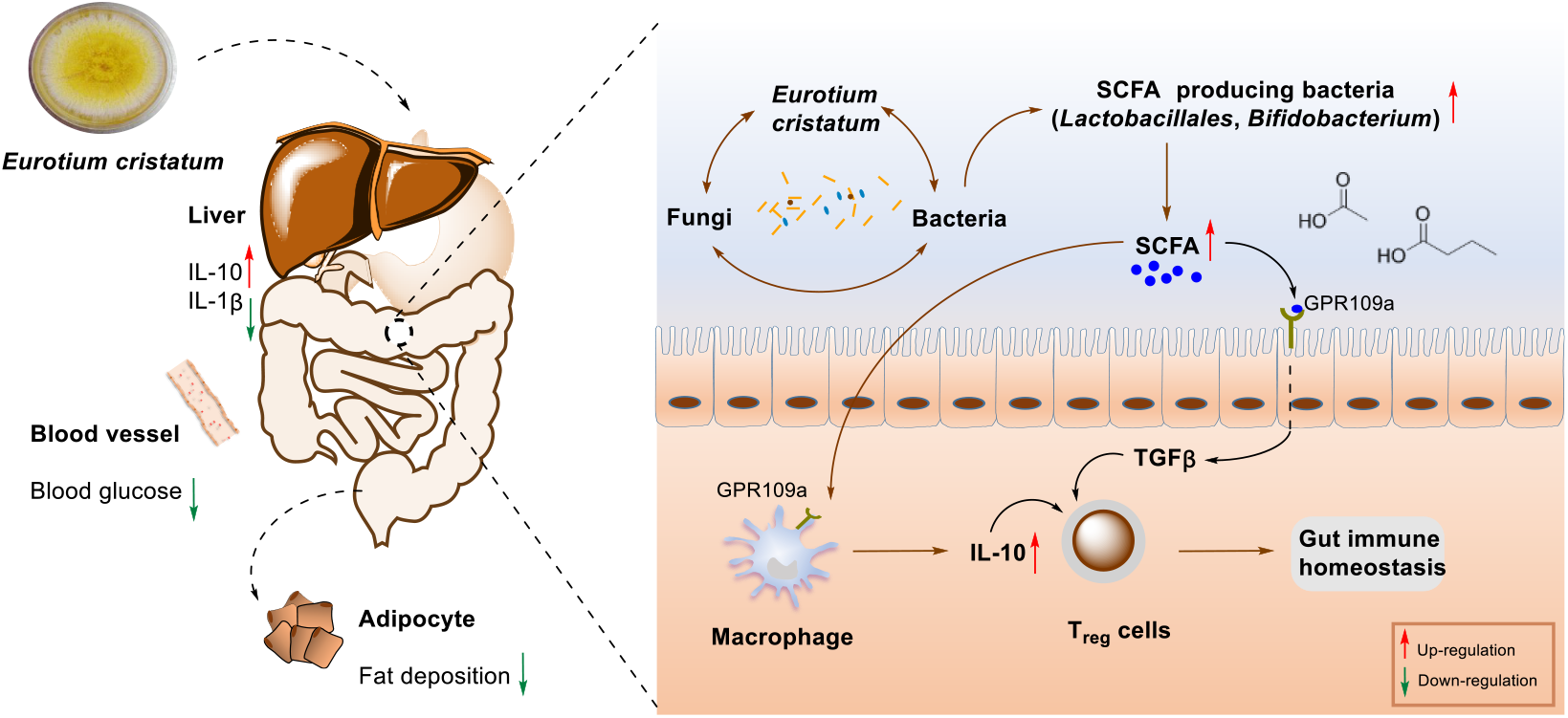
The proposed mechanism of dietary *E. cristatum* against HFD-induced obesity in mice. The interactions between dietary *E. cristatum* and gut microbiome may lead to the normalization of microbiota dysbiosis in the gut of HFD-fed mice. The increase of SCFAs-producing bacteria, including *Bifidobacterium* and *Lactobacillus*, leads to the production of more SCFAs in mice gut, especially acetate and butyrate. These changes lowered inflammatory cytokines and led to the observed anti-obese effects.

The gut microbiota of obese humans and animals have decreased ratio of Bacteroides-to-Firmicutes, and Firmicutes are crucial in obesity-related metabolic disorder^41^. Therefore the treatment of obese mice with *Myrciaria dubia*^42^, melatonin^43^ and other probiotics reversed this ratio and alleviated obesity. The consumption of FBT in HFD-fed mice could restore this lopsided microbiota, while supplementation of FFBT and *E. cristatum* were unable to reverse this unbalance. Probiotic bacteria, such as SCFAs-producing *Lactobacillus, Bifidobacterium* and *Leuconostoc* were shown previously to prevent or treat gastrointestinal disorders^44–46^, and to have anti-obesogenic or anti-diabetic potential^47–49^. Our study demonstrated that *E. cristatum* enhanced *Lactobacillus, Bifidobacterium* and *leuconostoc* in mice gut. *Akkermansia muciniphila* were shown to tighten gut barrier and reversed high-fat diet-induced obesity and improve glucose tolerance under high-fat diet conditions in conventional mice^15,50,51^. *A. muciniphila* in the *E. cristatum-treated* HFD-fed mice group also increased slightly. These results suggested that *E. cristatum* could modulate gut microbiota by increasing beneficial bacteria in mice gut. Although there is a strong correlation between gut fungi and host health, few studies were reported to manage metabolic disorder induced by high-fat diet by regulating gut fungi^33,34^. Our study also demonstrated that *E. cristatum* were able to modulate both fungi and bacteria composition (Fig. 3-4).

SCFAs, generated by the fermentation of intestinal microbiota^52,53,54^, are important to control chronic inflammatory diseases^55,56^ and maintain gut homeostasis^57^. The generation of a large amount of butyrate in the gut of *E. cristatum-treated* HFD-fed mice was unprecedented, which might contribute to the observed anti-obesity effects. Locally produced butyrate played essential role for the development of intestinal and systemic immune system. For example, butyrate may directly or indirectly induce T_reg_ differentiation, including IL-10 producing T_reg_ cells. It was consistent with the increased level of IL-10, and the decreased level of IL-1β in *E. cristatum-treated* HFD-fed mice (Fig. 2). Similar effects were also observed in FBT-treated HFD-fed mice.

In conclusion, our study strongly suggested that *E. cristatum* had significant anti-obesity effects in HFD-fed mice by modulating gut microbiota. Therefore, it may be used as a fungi probiotic to alleviate obesity, or other metabolic diseases, such as type II diabetes involving microbiota dysbiosis. Although Fuzhuan brick tea has already been consumed by humans for hundreds of years, the potential toxicity of the direct consumption of *E. cristatum* in humans needs to be established (Fig. S1). Since low dosage and short period of *E. cristatum* consumption may also have certain anti-obesity effects in HFD-fed mice (Fig. 7), *E. cristatum* may be used as a promising probiotic to alleviate obesity epidemic in the near future.

## Methods

### Preparation of *E. cristatum*, FBT and FFBT

*E. cristatum* CB10001 was isolated from Fuzhuan brick tea (FBT) (Jiuyang Processing Factory Co., Ltd., Yiyang, Hunan, China), which was confirmed to be *E. cristatum* based on its 18S rRNA sequence. *E. cristatum* CB10001 was grown in M40Y medium and its spores were harvested and stored in 20% glycerol and kept in –80 °C. The extracts for Fuzhuan brick tea (FBT) were prepared by brewing the tea with 85 °C deionized H_2_O, which was previously boiled. Briefly, Fuzhuan brick tea (8 g) was brewed in 1 L deionized H_2_O (85 °C) in a household teapot for 15 min. The filtered Fuzhuan brick tea (FFBT) was prepared by filtering the prepared FBT through a defecator (0.22 μm, thermo, USA).

### Survival of *E. cristatum* in elevated temperature

*E. cristatum* (10^4^ CFU) were suspended in 1 mL 20% glycerol and heated for 2 min, 5 min, 10 min under 50, 70, 80, 85, 90 and 100 °C, respectively. Then *E. cristatum* in each sample was serially diluted and spread onto M40Y agar plate. The plates were incubated at 30 °C for 48 h, and the resulting colonies were counted.

### Dosage

Typically an adult (60 kg) may consume 10 g of Fuzhuan brick tea per day. Therefore the dosage of a mouse to consume the Fuzhuan brick tea was set as 1.6 mg/g (1.6 mg/g = 10 g/60 kg × 10) Fuzhuan brick tea per gram of individual mouse weight, which was about 10 times of Fuzhuan brick tea a human might have consumed. Since Fuzhuan brick tea contains *E. cristatum* (2 × 10^5^ CFU/g), the dosage of *E. cristatum* per mouse was set at 10^3^ CFU/day first. To optimize the dosage of *E. cristatum*, 10^2^ or 10^4^ CFU/day/mouse of *E. cristatum* were also used.

### Animals

All animals in this study were purchased from Hunan Silaikejingda Experimental Animal Company Limited (Hunan, China). They were bred in the specific pathogen-free animal (SPF) facility of the Department of Laboratory Animals at Central South University (Changsha, Hunan, China) and kept under controlled light conditions (12 h light–dark cycle), with free access to water and diet.

Kunming mice (4-week, male) were randomly distributed into five groups, and each group had five mice. They had free access to chow diet containing 13.5% fat. The mice in normal chow diet group (NCD) had free access to drinking water. The mice in *E. cristatum* groups (10^4^ CFU/day) or (10^3^ CFU/day) only had access to drinking water containing different amount of *E. cristatum*. The mice in the Fuzhuan brick tea group (FBT) only had access to FBT as drinking water, while the mice in the filtered Fuzhuan brick tea group (FFBT) only had access to FFBT as drinking water. All the treatment groups, including *E. cristatum* (10^4^ CFU/day), *E. cristatum* (10^3^ CFU/day), FBT and FFBT were given at every other 4-weeks, until the end of 16^th^ week. The body weight of individual mouse was measured every other week and the feces of the mice were also collected every other week.

C57BL/6J mice (6-week, male) were randomly distributed into five groups, and each group contained five mice. They had free access to chow diet (13.5% fat) or high-fat diet (60% fat) (Trophic Animal Feed High-Tech Co. LTD, Jiangshu, China). *E. cristatum* (10^3^ CFU/day), FBT and FFBT were given to HFD-fed mice in drinking water for 7 weeks.

### Optimal dosage of *E. cristatum*

To identify the optimal dosage of *E. cristatum*, C57BL/6J mice (6-week, male) were randomly divided into 7 groups and each group had 10 mice. They had free access to chow diet (13.5% fat) or high-fat diet (60% fat). The 7 groups included: normal chow diet group (NCD), high-fat diet group (HFD), high dosage of *E. cristatum* group (10^4^ CFU/day, 8 weeks), middle dosage of *E. cristatum* group (10^3^ CFU/day, 8 weeks) and low dosage of *E. cristatum* group (10^2^ CFU/day, 8 weeks), as well as *E. cristatum* group (10^3^ CFU/day, 3 weeks) and *E. cristatum* group (10^3^ CFU/day, 5 weeks), in which the HFD-fed mice were given *E. cristatum* for only 3 or 5 weeks, respectively.

### Isolation and identification of *E. cristatum* from Kunming mice

The feces of Kunming mice (200 mg) were suspended in 1 mL sterile water. Then 100 μL supernatant was spread onto M40Y agar plates and cultured at 30°C for 48h. The 18S rDNA were PCR amplified from the isolated strain. The phylogenetic analysis of the 18S rRNA was performed using Mega 6 using the Neighbor-Joining method.

### Oral glucose tolerance test

Overnight-fasted mice were administered a 2 g kg^-1^ OGTT (20% glucose solution) by oral gavage. Their blood glucose was measured by tail bleeding at time 0, 15, 30, 60, 90 and 120 min after oral gavage, using a glucose meter (Sannuo Biosensor Co., Ltd, Hunan, China).

### Cytokine measurements

IL-1β, IL-10 protein levels were measured using commercial ELISA kits (eBioscience, USA).

### Gut microbiota analysis

Mice feces were stored at –80 °C, and the total DNAs were extracted using a stool DNA extraction kit (Omega, USA). For each caecal stool sample, the 16S rRNA gene comprising V4 regions were amplified, using a forward primer 515F (5’-GTGCCAGCMGCCGCGGTAA-3’) and a reverse primer 806R (5’-GGACTACHVGGGTWTCTAAT-3’). Both primers also contained a unique 12-base barcode to tag each PCR product. The ITS2 region of fungi in the caecal stool samples were amplied using a forward primer ITS7F (5’-GTGARTCATCGARTCTTTG-3’) and a reverse primer ITS4R (5’-TCCTCCGCTTATTGATATGC-3’), which also contained a unique 12-base barcode to tag each PCR product. The high throughput sequencing was carried out using the Illumina MiSeq platform to generate 2 × 250 bp paired-end reads. The raw reads were quality filtered. The primers and barcode in each read were trimmed using QIIME^58^ software package. The resulting sequences were then assigned to operational taxonomic units (OTUs) with 97% identity, followed by the selection of representative sequences. The alpha and beta diversity were determined using a UniFrac analysis and a constrained principal coordinate analysis (CPCoA). The taxonomic abundance and characterize differences between groups were estimated by Linear discriminant analysis of the effect size (LEfSe). Phylogenetic investigation of communities by reconstruction of unobserved states (PICRUSt)^39^ was performed to identify functional genes in the sampled microbial community on the basis of the data in the KEGG pathway database.

### SCFA analysis

SCFAs were similarly analyzed as described before^59^. Briefly, 50 mg fecal sample was blended with 500 μL acetonitrile and acidified with 5% (v/v) H**2**SO**4**. Then the mixture was ultrasonically extracted for 10 min and centrifuged at 15000 rpm for 10 min. The supernatant was filtered through a 0.22 μm filter. SCFAs were measured using a gas chromatograph-mass spectrometer (GCMS-QP2010 Ultra system, Shimadzu Corporation, Japan), which was equipped with an Rtx-Wax column (30 m × 0.25 μm × 0.25 μm). The carrier gas was helium (flow 2 mL min^-1^, split ratio 10: 1, the volume of sampling 1 μL). The injection and ionization temperature was set at 200 °C. The standard curves of acetate, propionate and butyrate were established and the concentrations of these SCFAs (mmol/kg mouse feces) were calculated according to the standard curve.

### Statistics

Statistical analysis was performed using GraphPad Prism 6.01 (GraphPad Software, Inc., San Diego, CA) unless otherwise specified. Data obtained from experiments are shown as means ± s.e.m. Differences in body weight and OGTT were analyzed by the unpaired two-tailed Student’s t-tests, and those with more than two groups were assessed with one–way ANOVA followed by Newman–Keuls *post hoc* tests. In the figures, the data with different superscript letters are significantly different based on post hoc ANOVA statistical analysis. Graph bars marked with different letters on top represented statistically significant results (P<0.05) based on Newman–Keuls post hoc one–way ANOVA analysis, whereas bars labeled with the same letter correspond to results that show no statistically significant differences. In the case, whereas two letters are present on top of the bar, each letter should be compared separately with the letters of other bars to determine whether the results show statistically significant differences.

### Accession codes

Both sequencing data for the 16S rRNA and ITS sequences have been deposited in the NCBI’s Sequence Read Archive database under BioProject ID No PRJNA506283. The 18S rRNA of *E. cristaum* CB10001 has been deposited in GenBank (MK371789).

## Acknowledgements

The authors thank the Center for Advanced Research for the GCMS and Minerals processing and bioengineering for Illumina MiSeq platform in Central South University. This work was supported in parts by NSFC grant 81473124 (to Y. H.), the Chinese Ministry of Education 111 Project B0803420 (to Y. D.).

## Author contributions

Y.H. and Y.D. conceived the project; Y.H., Y.D. and D.K. designed the experiments; D.K. and M.S. performed the experiments; Y.H., and D.K. analyzed the results and wrote the manuscript with inputs from all co-authors.

## Competing interests

The authors declare no competing financial interests.

## Materials & Correspondence

All the correspondence and material requests should be addressed to (Y.H.) or (Y.D.).

## Animal protocols

All animal protocols were approved by the Animal Care and Use Committee of Central South University. All experimental procedures complied with the Guide for the Care and Use of Laboratory Animals (1996).

## Additional information

Supplementary information is available online.

## Competing interests

There are no conflicts to declare.

