## Supporting Information for "*Eurotium cristatum*, a new fungi probiotic from Fuzhuan brick tea, alleviated obesity in mice by modulating gut microbiota"

### Table of Contents


**Tables S1** Summary of pilot dietary intervention of type 2 diabetes or metabolic syndrome using Fuzhuan brick tea…………………………………..……………..……......**S3**

**Fig. S1.** Effects of heating temperature on the survival of *E. cristatum*.......................................................................................................**S4**

**Fig. S2.** *E. cristatum* could survive in mouse intestine...................................................**S5**

**Fig. S3.** Diet and water consumption of mice in the first week of treatment for C57BL/6J mice fed with normal chow diet or high fat diet.........................................................................................................**S6**

**Fig. S4.** Alpha diversity analysis of mice microbiota for C57BL/6J mice.......................................................................................................**S7**

**Fig. S5.** *E. cristatum,* FBT and FFBT beneficially altered the gut metabolism of high-fat fed mice.......................................................................................................**S8**

**Table S1 | Summary of pilot dietary intervention of type 2 diabetes or metabolic syndrome using Fuzhuan brick tea**[1-2]. In reference 1: the patients in the control treatment group were orally given metformin (0.5 g × 2/day) and the traditional Chinese medicine Danshen pill (10 capsules × 3/day)；The patients in the treatment group were orally given mulberry black tea 30 g (mulberry leaf 1: 5 Fuzhuan brick tea), as well as metformin (0.5 g × 2/day) and the traditional Chinese medicine Danshen pill (10 capsules × 3 /day). The treatments lasted for 12 weeks. * *P*<0.05: there were significant differences before treatment and after treatment after 12 weeks; # *P*<0.05: there were significant differences between the treatment and control groups after 12 weeks. In reference 2: the patients were given Fuzhuan brick tea (15 g/day) for 1 month. * *P*<0.05, ** *P*<0.01: there were significant differences before treatment and after treatment after 1 months. FBG, fasting glucose; OGTT, oral glucose tolerance test; TC, total cholesterol; TG, triglyceride; HbA1c, hemoglobin a1c.

| **Index** | **Before treatment** | | **After treatment** | | **Number of patients** | **References** |
|---|---|---|---|---|---|---|
| | Treatment group | Control group | Treatment group | Control group | | |
| FBG (mmol/L) | 8.81±1.87 | 8.66±1.30 | 6.47±1.09*# | 7.31±1.42* | Control group: 20<br>Treatment group: 20 | 1 |
| OGTT 2h (mmol/L) | 10.58±1.62 | 10.32±2.09 | 8.21±1.34*# | 9.13±1.50* | | |
| HbA1c (%) | 7.11±1.00 | 7.38±1.47 | 6.01±0.58*# | 6.42±0.65* | | |
| LDL-C (mmol/L) | 3.32±0.37 | 3.44±0.51 | 2.49±0.39*# | 2.77±0.39* | | |
| HDL-C (mmol/L) | 0.87±0.13 | 0.85±0.14 | 1.17±0.17*# | 1.06±0.13* | | |
| Blood uric acid (μmol/L) | 404±39 | 415±42 | 347±45*# | 381±52* | | |
| FBG (mmol/L) | 10.04±2.95 | | 8.06±1.78* | | Treatment group: 45 | 2 |
| TC | 6.7±0.79 | | 5.75±0.78* | | | |
| TG | 3.3±1.43 | | 1.86±1.12** | | | |
| HDL-C (mmol/L) | 1.3±0.40 | | 1.34±0.44 | | | |
| LDL-C (mmol/L) | 4.2±0.73 | | 3.71±0.60* | | | |

* *P*<0.05 ** *P*<0.01

**Fig S1 | Effects of heating temperature on the survival of *E. cristatum*.** $10^4$ colony forming units (CFU) of *E. cristatum* were suspended in 1 mL 20% glycerol and heated for 2 min, 5 min, 10 min respectively. The CFU of *E. cristatum* were counted on agar plates containing M40Y medium. There were 3 replicates for every temperature tested.

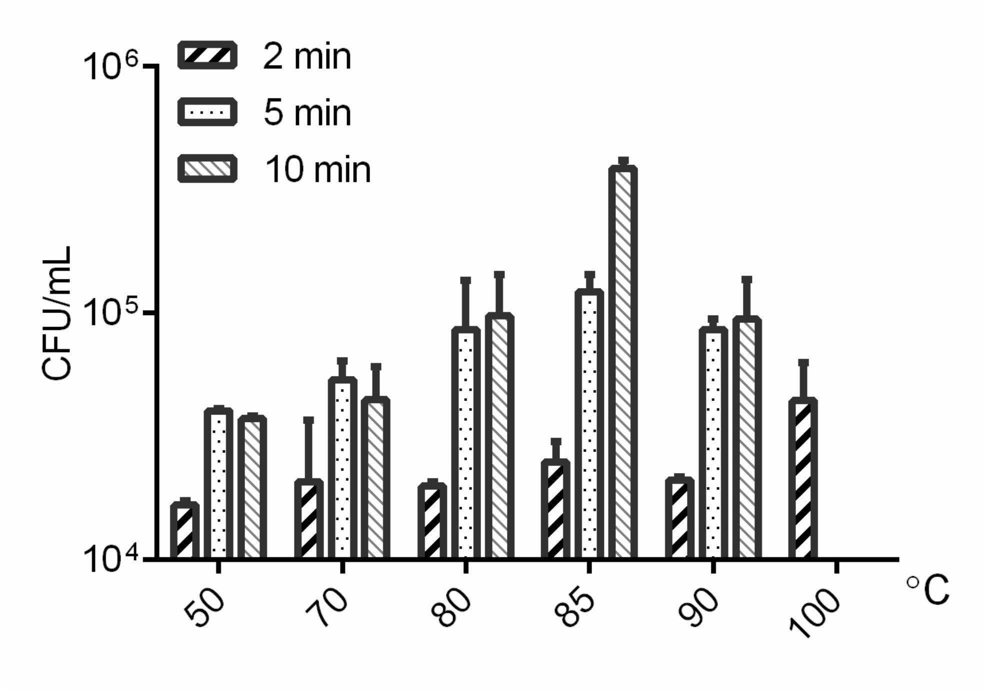

**Fig S2 | *E. cristatum* could survive in mouse intestine. a**, The timeline for mouse experiments. **b**, The mice body weights of different groups. NCD, normal chow diet group; FBT, normal chow diet with Fuzhuan brick tea; FFBT, normal chow diet with filtered Fuzhuan brick tea; *E. cristatum* ($10^4$ CFU/day) and *E. cristatum* ($10^3$ CFU/day): normal chow diet with the indicated amount of *E. cristatum* in the drinking water. There were 5 mice per group. **c**, the growth of *E. cristatum* from the feces of NCD group (left) and *E. cristatum* ($10^4$ CFU/day) (right) mice at the end of 6th week. **d**, Phylogenetic analysis of the 18S rRNA of *E. cristatum* CB10001 isolated from mice feces using Mega 6 and the Neighbor-Joining method. *Chaetosartorya cremea*: AB002074.1; *Fennellia flavipes*: AB008400.1; *Neosartorya fischeri*: U21299.1; *Eurotium amstelodami*: AB002076.1; *Aspergillus terreus*: AB008409.1; *Aspergillus penicillioides*: AB002078.1; *Aspergillus penicillioides* strain 481: DQ985958.1; *Eurotium cristatum*: JN986762.1.

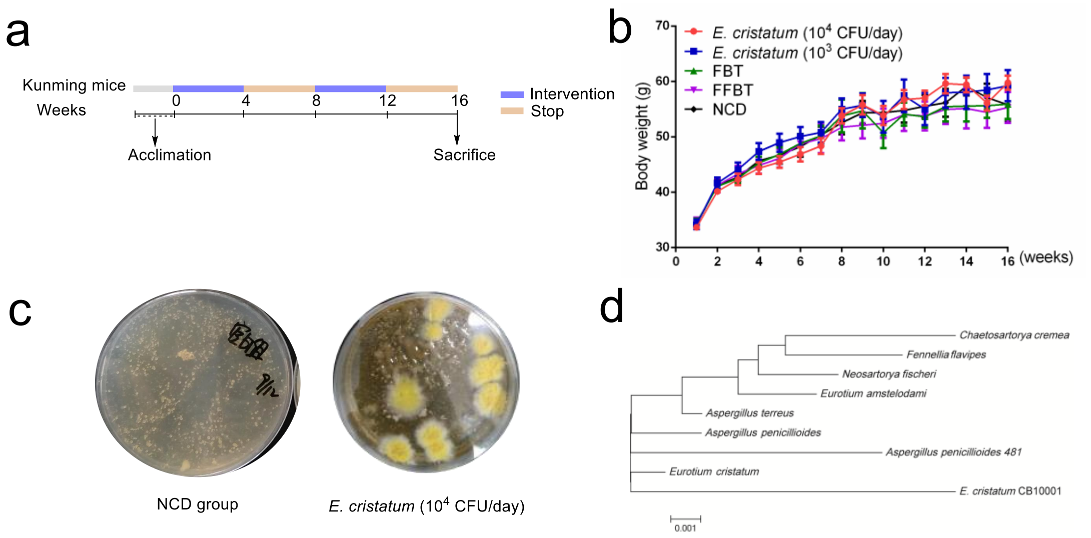

**Fig S3** | **Diet and water consumption of mice in the first week of treatment for C57BL/6J mice fed with normal chow diet or high-fat diet. a**, Diet consumption of each group. **b**, Water consumption of each group. Data are expressed as mean ± s.e.m. Graph bars in **a** and **b** marked with different letters on top represent statistically significant results (P<0.05) based on Newman-Keuls *post hoc* one-way ANOVA analysis, whereas bars labelled with the same letter correspond to results that show no statistically significant differences. In the case whereas two letters are present on top of the bar, each letter should be compared separately with the letters of other bars to determine whether the results show statistically significant differences. NCD, normal chow diet group; HFD, high-fat diet group; FBT, high-fat diet with consumption of Fuzhuan brick tea; FFBT, high-fat diet with the consumption of filtered Fuzhuan brick tea. *E. cristatum*, high fat diet with consumption of *E. cristatum* ($10^3$ CFU/day). There were 5 mice per group.

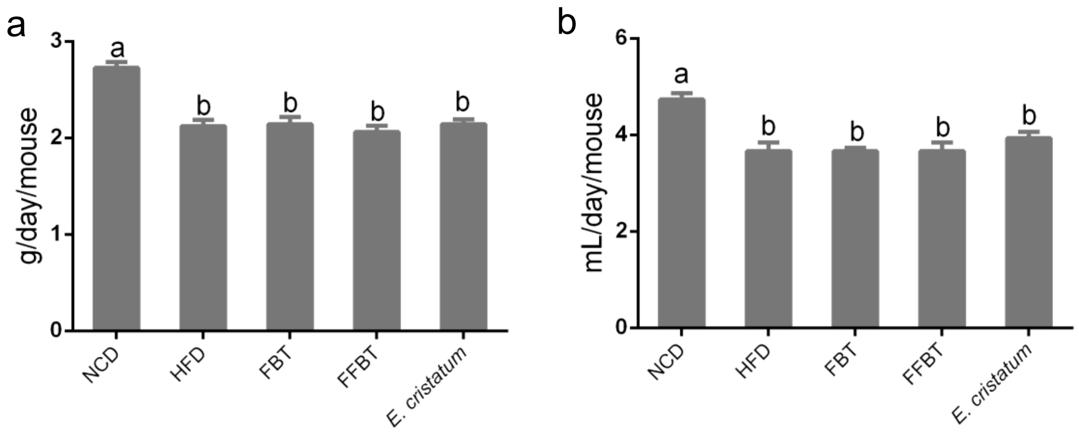

**Fig S4 | Alpha diversity analysis of mice microbiota for C57BL/6J mice. a**, Rarefaction (chao1) analysis of gut fungi in each treatment group. **b**, Rarefaction (observed OTUs) analysis of gut fungi in each treatment group. **c**, Rarefaction (chao1) analysis of gut bacteria in each treatment group. **d**, Rarefaction (observed OTUs) analysis of gut bacteria in each treatment group. NCD, normal chow diet group; HFD, high-fat diet group; FBT, high-fat diet with consumption of Fuzhuan brick tea; FFBT, high-fat diet with the consumption of filtered Fuzhuan brick tea. *E. cristatum*, high fat diet with consumption of *E. cristatum* ($10^3$ CFU/day). There were 5 mice per group.

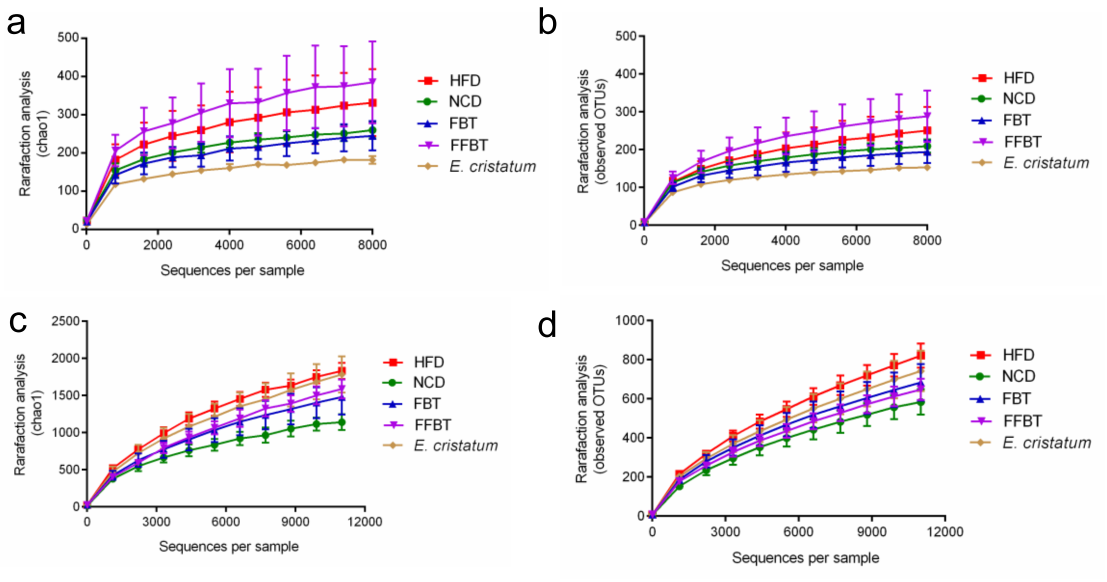

**Fig S5 | *E. cristatum*, FBT and FFBT beneficially altered the gut metabolism of high-fat fed mice.** PICRUSt analysis showed the relative abundance of predicted microbial genes related to metabolism for **a**, *E. cristatum* and HFD, **b**, FBT and HFD, **c**, FFBT and HFD based on Welch's *t* test (*P*<0.05). The colored circles represent 95% confidence intervals calculated using Welch's inverted method. NCD, normal chow diet group; HFD, high-fat diet group; FBT, high-fat diet with consumption of Fuzhuan brick tea; FFBT, high-fat diet with the consumption of filtered Fuzhuan brick tea. *E. cristatum*, high-fat diet with consumption of *E. cristatum* ($10^3$ CFU/day). There were 5 mice per group.

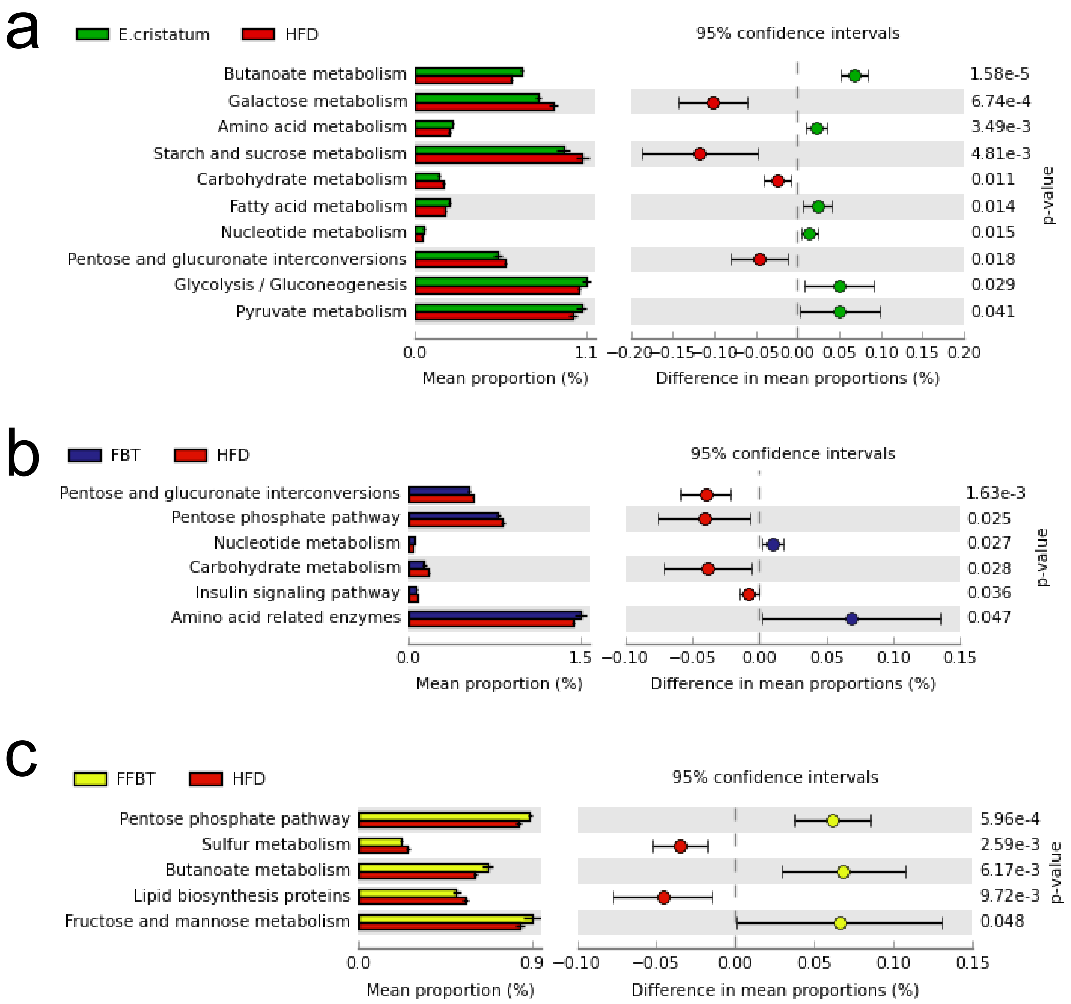
